# Montane Central Appalachian Forests Provide Refuge for the Critically Endangered Rusty Patched Bumble Bee (*Bombus affinis*)

**DOI:** 10.1101/2023.10.19.563135

**Authors:** Mark J. Hepner, Ellison Orcutt, Kyle Price, Karen Goodell, T’ai Roulston, Robert P. Jean, Rodney T. Richardson

**Affiliations:** Metamorphic Ecological Research and Consulting, LLC, Alonzaville, VA 22644; Division of Natural Heritage, Virginia Department of Conservation and Recreation, Richmond, VA 23219; Environmental Solutions & Innovations, Inc. Indianapolis, IN 46241; Department of Evolution, Ecology and Organismal Biology, The Ohio State University, Newark, OH 43055; Department of Environmental Sciences, University of Virginia, Charlottesville, VA 22908; Appalachian Laboratory, University of Maryland Center for Environmental Science, Frostburg, MD 21532

**Keywords:** Pollinator conservation, random forest, climate velocity, *Bombus terricola*

## Abstract

The mountains of Central Appalachia are rich with environmental variance and host a wide variety of community types and diverse flora and fauna. The once common Rusty-Patched Bumble Bee (RPBB, *Bombus affinis*) has experienced widespread declines and was believed to have been extirpated throughout the Lower Midwest, Northeast and Appalachian regions of the United States (U.S.). We document the occurrence and environmental associations of a contemporary population within Central Appalachia using a dataset of 274 observations spanning nine years and over 2,000 surveys. We show that Appalachian RPBB are strongly associated with high elevation, heavily forested landscapes, especially those with West to Northwest facing aspects. Measures of forest species composition are also associated with RPBB observations. While only 38 percent of surveys occurred on U.S. National Forest lands, 84 percent of observations occurred in these areas, suggesting distinct forest habitat conditions associated with U.S. Forest Service lands play a role in the persistence of this species. The Appalachian region is rugged and difficult to systematically survey, and our analysis represents the first assessment of the species presence and habitat associations within the region. Appalachian RPBB populations are likely geographically and genetically isolated from Upper Midwest populations and additional research is needed to prioritize future conservation efforts across the current and potential range of the species.

## Introduction

The mountains of Appalachia encompass diverse and heavily forested temperate ecoregions (Pickering et al., 2002; Stein et al., 2000) characterized by gradients in elevation, aspect and soil type that in turn drive heterogeneity in ecosystems and community types (Fekedulegn et al., 2003, 2004; Whittaker, 1956). This environmental variability, coupled with differing land management approaches, including diverse timbering practices, controlled burns, and pastoral agriculture have contributed to a mosaic of habitats and legacy effects at the landscape scale (Arthur et al., 2015; Gragson & Bolstad, 2006; Silver et al., 2013; Thomas-Van Gundy et al., 2014; Wyatt & Silman, 2010). Relatively high species richness and habitat connectivity further contribute to the considerable conservation value found within this region (Belote et al., 2016; McKinley et al., 2019).

Historical survey data for bumble bees (*Bombus*) in the Appalachian Mountains indicate that the Rusty Patched Bumble Bee (RPBB, *Bombus affinis*) once occurred from Georgia, U.S. to New Brunswick, Canada (Klymko & Sabine, 2015; Mitchell, 1962). Following sharp population declines starting in the 1990s (Colla & Packer, 2008; Grixti et al., 2009), RPBB was believed to have been extirpated from all areas of the species’ original range east of Illinois (Cameron et al., 2011). Range contraction of the RPBB is not unique; worldwide, about one third of bumble bee species are recognized as declining, with clear phylogenetic patterns regarding which taxonomic groups are most affected (Arbetman et al., 2017). In North America, the subgenera *Thoracobombus* and *Bombus* appear particularly vulnerable, with all eight North American species considered to be in decline (Cameron et al., 2011; Arbetman et al., 2017; Jacobson et al., 2018). The causes of bumble bee decline are not clear, but one common trait across declining species is a disproportionately high occurrence of infections by the microsporidian *Vairimorpha bombi* (Cameron et al., 2011), which can cause colony failure (Otti & Schmid-Hempel, 2007). Other possible stressors to bumble bees include drought, climate change, and neonicotinoid pesticide exposure (Soroye et al., 2020; Janousek et al., 2023). Risks associated with these stressors appear particularly pronounced for members of the subgenus *Bombus*, which includes multiple imperiled species such as RPBB and the western bumble bee (*B. occidentalis*), though it remains untested whether these species are more sensitive to climate change and pesticides relative to other bumble bees.

Sightings of RPBB in the Mid-Atlantic U.S. over the last two decades suggested that the species persisted in the region, but initial data were too sparse to discern patterns of occurrence relative to habitat characteristics. The first contemporary sighting in Appalachia occurred in 2014, when a single RPBB female was caught in a blue vane trap survey at an elevation of 213 m in the foothills of the Blue Ridge Mountains in Fauquier County, Virginia. Three years later, in the same year RPBB was listed under the U.S. Endangered Species Act, a female was netted from flowers of *Rhododendron* sp. during a survey in Bath County, VA and a male was netted from *Helianthus* sp. in Mineral County, West Virginia (WV) (Sam Droege; personal communication). Over five subsequent years, as search efforts increased (Table 1), RPBB were observed with increasing annual frequencies in the high elevation forests of the region, predominantly in the U.S. Forest Service (USFS) Monongahela and George Washington and Jefferson National Forests. Meanwhile, no additional observations have been recorded at the original 2014 site or anywhere below 450 m of elevation. These sightings suggest that cool temperatures, forest cover, or factors associated with these variables favor the persistence of eastern U.S. RPBB populations, although this interpretation is complicated by the absence of RPBB in other mountainous regions of its former range, such as in New England.

**Table 1:** Summary of survey effort and RPBB detections by year.

|  | Number of Surveys | Survey Period (DOY) | RPBB Positive Surveys | Teams Surveying |
| --- | --- | --- | --- | --- |
| 2014 | 115 | 127-241 | 1 | 1 |
| 2015 | 27 | 163-220 | 0 | 1 |
| 2016 | 67 | 161-219 | 0 | 1 |
| 2017 | 100 | 157-223 | 1 | 2 |
| 2018 | 288 | 159-283 | 8 | 3 |
| 2019 | 366 | 144-284 | 31 | 4 |
| 2020 | 118 | 146-288 | 10 | 2 |
| 2021 | 452 | 81-262 | 38 | 3 |
| 2022 | 610 | 135-252 | 59 | 5 |

In this study, we report the spatial distribution and habitat characteristics associated with recent RPBB sightings within Appalachia. We aggregated survey efforts across multiple independent teams to quantify associations between Appalachian RPBB detections and habitat features in order to gain insight into the biology of the species and guide future survey efforts. We hypothesize contemporary Appalachian RPBB observations are positively associated with landscape-scale measures including: 1) elevation, 2) cooler and more mesic northwest facing aspects, 3) proportion of the landscape that is forest, 4) land ownership by USFS, and 5) basal area of tree species considered to provide foraging resources for bumble bees. Our work informs expectations about the future persistence of Appalachian RPBB in response to ongoing anthropogenic forces and provides a predictive framework for future surveys of the species outside of areas which are currently known to be inhabited by the species.

## Methods

### Field Surveys and Data Collection

Survey data were collected by five independent research teams over the course of nine years, 2014 to 2022, throughout the Central Appalachian Mountains and adjacent piedmont within VA, WV, Pennsylvania (PA), and Maryland (MD). Though most of the data (i.e. greater than 1,500 surveys) consisted of standardized 10-minute searches, sampling effort varied across the remaining surveys, and we do not attempt to account for this variance in effort during statistical analysis. Instead, we rely on a large sample size as well as the decentralized nature of our data collection. To minimize pseudo-replication, we combined data from surveys that occurred on the same date within the same 250 m^2^ grid cell (based on FIA grid cells; see next section). On any given survey date where a grid cell was surveyed multiple times, surveys were collapsed and recorded as a single replicate, and marked positive if RPBB was observed in any individual survey. Upon documenting RPBB within an area, sampling is limited to the time period approved by the U.S. Fish and Wildlife Service (June – August), which is intended to minimize the potential impacts of surveys on queen bumble bees (USFWS 2019). Therefore, we rarely surveyed during the early and later periods of the historical flight season of RPBB (April -- October), and most survey effort was conducted when the species is expected to be relatively abundant.

### Habitat covariates

We quantified RPBB environmental associations using a comprehensive set of covariates that characterized variation in climate, topography, land cover, and forest tree composition within our study region. For climate, we extracted average monthly maximum and minimum temperature and monthly sum of precipitation for the period 2011-2021 within the study region using the *get_daymet* function in the FedData package (Bocinsky 2022). We then used the *biovars* function in the dismo package (Hijmans et al. 2022) to calculate the suite of 19 bioclimatic variables from the 10-year time series of monthly Daymet data, which are provided at 1 x 1 km resolution (Thornton et al., 2022). To characterize topography, we used the *get_elevation_raster* function in the elevatr package (Hollister 2021) to download elevation data from the Open Topography global dataset. Within the *get_elevation_raster* function, we set the z parameter (zoom level) to 10, which resulted in a raster resolution of approximately 100 m^2^. We then used the terrain function in the raster package (Hijmans 2023) to calculate aspect, slope, terrain ruggedness Index (TRI), and roughness. Land cover was characterized using the 2019 National Land Cover Dataset (NLCD; Dewitz et al., 2021) and downloaded for our study region using the *get_nlcd* function in the FedData package (Bocinsky 2022). Lastly, we estimated tree species composition and abundance using a 250-m resolution gridded product based on USFS

Forest Inventory and Analysis (FIA) data, MODIS remote sensing of phenology, and environmental parameters (Wilson et al. 2012). These data comprise an individual 250-m resolution raster of estimated basal area for each tree species measured by the FIA program. A moving window analysis was used to calculate summary statistics for each of our variables within a radius of 500, 1000, and 3000m of each 250-m grid cell containing a survey. Within these windows, we calculated averages of climate and topography covariates, proportion of each land cover type, and sum of basal area of each tree species. A full list of covariates included in this dataset is shown in Supplemental Table S1. During all statistical analysis, three additional variables were calculated: *Forest* (the sum of NLCD mixed, deciduous and evergreen forest categories), *Ag* (the sum of NLCD categories Pasture or Hay and Cultivated Crops) and *Urban* (the sum of NLCD developed open space as well as low, medium and highly developed categories).

### Statistical Analysis

We first generated a montane-centric generalized linear model (GLM) to summarize the biophysical associations of the species with regard to features such as *Elevation* and *Aspect*, proxy measures indirectly associated with other climatic and biological features. With this montane model, we used model selection to substitute *Elevation* and *Aspect* with the bioclimatic variables, creating a second model that could be generalized across broader spatial scales. We then probed further into an observed association between RPBB and forest composition, inferred from the montane-centric model, incorporating the basal areas of dominant tree genera, those with a cumulative basal area greater than or equal to one million square feet per acre across all survey sites. All models were assessed by calculation of area under the receiver operating characteristic curve (AUC) using the average of 100 bootstrapped out-of-bag (OOB) estimates on 20 percent hold-out samples, calculation of McFadden’s pseudo-*R^2^*, and estimation of variance inflation factors (VIFs) for all predictor variables, excluding day of year (*Day*) which was modeled as a quadratic effect. Notably, the models presented here do not include the initial specimen caught in a blue vane trap in 2014 because it is a temporal outlier with the lowest average elevation, lowest percent forest cover and greatest *BIO8* estimate of all contemporary sightings (see Results selection below).

*Montane Generalized Linear Model* – Biophysical associates of RPBB presence were inferred using GLMs. We first determined the most appropriate spatial scale of analysis by regressing RPBB presence-absence outcomes against *Elevation* and *Forest*, with linear and quadratic effects for *Day*, at each of the three scales listed previously. The 3000 m radius scale was strongly supported relative to a 1000 m (ΔWAIC = +6.2) or 500 m scale (ΔWAIC = +8.4) and all variables were strongly associated with RPBB detection in this model (*P* < 0.001 for all coefficients). Using the 3000 m scale, we constructed a starting model with features agreed upon across coauthors. The starting model included linear and quadratic effects for *Day* and linear effects for *Elevation*, *Forest*, *Aspect* and basal area of bee-friendly tree species (*BTBA*, species list provided in Supplemental Table S2) within a 3000 m radius, as well as distance from nearest U.S. National Forest land (*NF*). Though we intended to conduct reverse-stepwise model selection, all terms were significantly associated with RPBB detection (*P* < 0.020) in this initial model, and therefore retained.

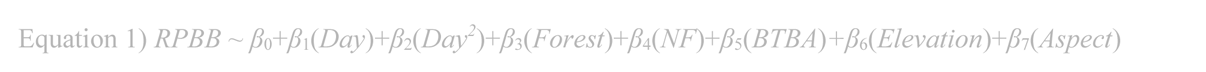

*Bioclimatic Generalized Linear Model* – After constructing the montane GLM, we substituted *Elevation* and *Aspect* with bioclimatic variables to create a more broadly generalizable model. We evaluated WAIC-based model fit on a set of models, each of which included the first five terms of Equation 1 and a linear effect term for one of the 19 bioclimatic variables. The bioclimatic variables from the five models with the lowest WAIC were selected for further review. Upon evaluating collinearities across these five variables, *BIO1* and *BIO10* were highly collinear (OLS *R^2^* = 0.981) and *BIO1* was excluded from further analysis due to a relatively higher WAIC score. Reverse-stepwise selection was implemented on the remaining four variables, where the starting model contained the first five terms of Equation 1 along with linear terms for *BIO8*, *BIO9*, *BIO10* and *BIO18*. Removal of the least significant of all non-significant (*α* = 0.05) features in each successive selection round resulted in a final model (Equation 2) which included *BIO8* (mean temperature of wettest quarter) and *BIO9* (mean temperature of driest quarter). This final model represented a strong improvement in fit relative to the montane model (ΔWAIC = -72.4), exhibited strong discriminatory power (ROC AUC = 0.919) and explained greater variance in the data relative to the original montane model (McFadden’s pseudo-*R^2^* = 0.379 relative to pseudo-*R^2^* = 0.311).

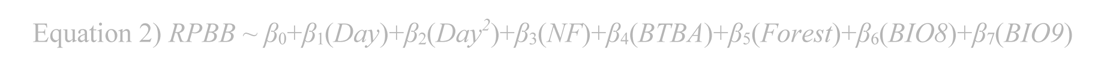

*Extended Forest Composition Modeling* – Given the unexpectedly strong negative association between RPBB and *BTBA*, we performed an additional round of model selection, incorporating the basal areas of two dominant tree genera, oak (*Quercus*) and pine (*Pinus*). These represent the two most abundant genera, estimated at a 3000 m scale, that were not included within the list of *BTBA* taxa. Basal areas were estimated for each site using the FIA Database and estimates from all congeneric species were summed to obtain genus-level basal area for each site. Reverse stepwise binomial regression was performed, starting with an initial model that included a quadratic effect for *Day*, linear terms for *BIO8, BIO9, Forest, NF* and *BTBA*, as well as linear terms for the basal areas of *Quercus* and *Pinus*. During each round of selection, the least significant of all non-significant terms was removed to create a reduced model, which ultimately resulted in the removal of the *BIO9, Forest* and *BTBA* terms.

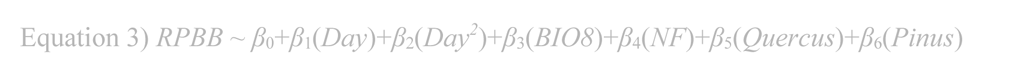

*Evaluating Influence of Agriculture and Urban Development* – The potential influence of agriculture and human development were assessed using reverse-stepwise selection. A starting model was created by combining two terms, *Ag* and *Urban*, with all terms from Equation 2. Removal of the least significant of all non-significant (*α* = 0.05) features in each successive selection round resulted in the exclusion of both *Ag* and *Urban*, resulting in reduction back to Equation 2. Alongside stepwise-selection, WAIC estimates across models provided additional support for the final reduced model relative to any competing specification (ΔWAIC ≤ -3.7).

*Evaluating Congruence between GLM Results and Random Forest Regression* - In addition to exploring our data with GLMs, we ran a set of random forest regressions to evaluate the congruence of trends between these two statistical approaches. During random forest regression, RPBB survey presence and absence outcomes were regressed against habitat covariates (measured at a 3 km radius scale) including landscape composition and topography measures, bioclimatic variables and the *Day* of the survey. Two weighted models and a third unweighted model were run to evaluate the robustness of results at multiple levels of down-weighting survey absences, which were about 20 times more prevalent in the data relative to detections. For the weighted models, weights were created such that the cumulative weight of all negative surveys equaled 1x and 5x the weight of all positive surveys. During surveys, RPBB negative effort is likely associated with greater uncertainty relative to positive survey effort. Thus, some readers may prefer to view a more conservative interpretation of effects from down-weighting negative surveys.

## Results

Across all five research teams, the full dataset included 2,143 surveys across 1,663 FIA grid cells. In total, 274 RPBB were observed across 147 surveys of 121 unique FIA grid cells. With regard to caste, 151 of these observations were female workers, while 111 were males and two were queens. Caste was undetermined for an additional 10 on-the-wing observations. These observations make up 87 percent of all contemporary RPBB sightings within Central Appalachia from 2014 to 2022, according to USFWS estimates (Tamara Smith, personal communication). RPBB were primarily observed in heavily forested, high-elevation sites within VA and WV, but three observations occurred in similar habitat in western MD. The lowest elevation landscapes in which detections occurred were approximately 1,800 feet in average elevation at the 3 km radius scale (Supplemental Figure S1). Female RPBB were predominantly observed on native flora within the families Lamiaceae (*Monarda*, *Pycnanthemum*), Asteraceae (*Solidago*, *Helianthus*, *Eutrochium*, *Verbesina*), Ranunculaceae (*Actaea, Thalictrum*) and Hydrangeaceae (*Hydrangea*). Additional native (*Rhododendron*, *Rubus*, *Aralia*, *Clematis*) and non-native (*Hibiscus syriacus*, *Cirsium vulgare*, *Carduus acanthoides*, *Nepeta, Trifolium pratense*) floral hosts were observed (Figure 1). Males were commonly observed on the same native flora, though a large proportion of sightings occurred on non-native plants (e.g. *Carduus acanthoides*, *Centaurea stoebe* and *Melilotus alba*).

**Figure 1:**
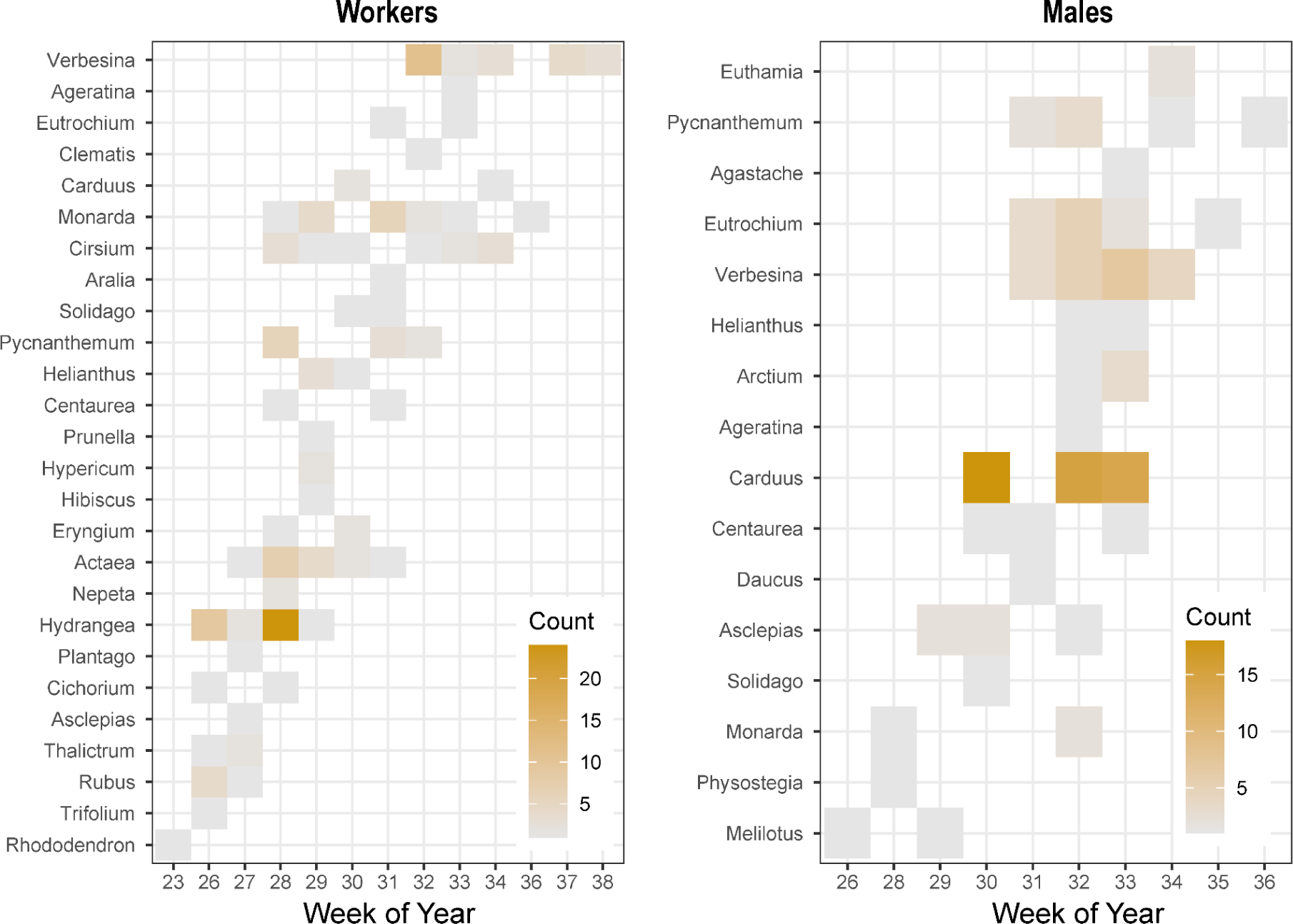
Counts of RPBB forage plant observations by genus and week of year, separated for worker and male castes.

An initial montane-centric model (Equation 1; Figure 2) relating RPBB presence and absence to survey and site-level attributes resulted in strong goodness-of-fit measures (McFadden’s pseudo-*R^2^* = 0.311, ROC AUC = 0.890) and significant effect estimates for all model terms. A strong non-linear effect for *Day* was observed (*β_Day_* = 5.0×10^-1^, *β_Day_^2^* = - 1.1×10^-3^, *P* < 0.001 for both terms) along with positive associations for *Elevation* (*β* = 1.3×10^-3^, *P* < 0.001), *Aspect* (*β* = 1.5×10^-2^, *P* = 0.017) and *Forest* (*β* = 4.6, *P* < 0.001). Negative effects were found for distance from U.S. National Forest land (*NF*; *β* = -4.3×10^-5^, *P* = 0.010) and the basal area of bee-friendly flowering tree species (*BTBA*; *β* = -2.3×10^-4^, *P* < 0.001). Minimal multi-collinearity was observed between linear predictors (VIF ≤ 2.03). Importantly, while *Day* was a strong predictor, a considerable fraction of the variation in RPBB survey outcomes was explained by habitat covariates as evidenced by a McFadden’s pseudo-*R^2^* of 0.228 when calculated using a null model which included a quadratic effect for *Day*.

**Figure 2:**
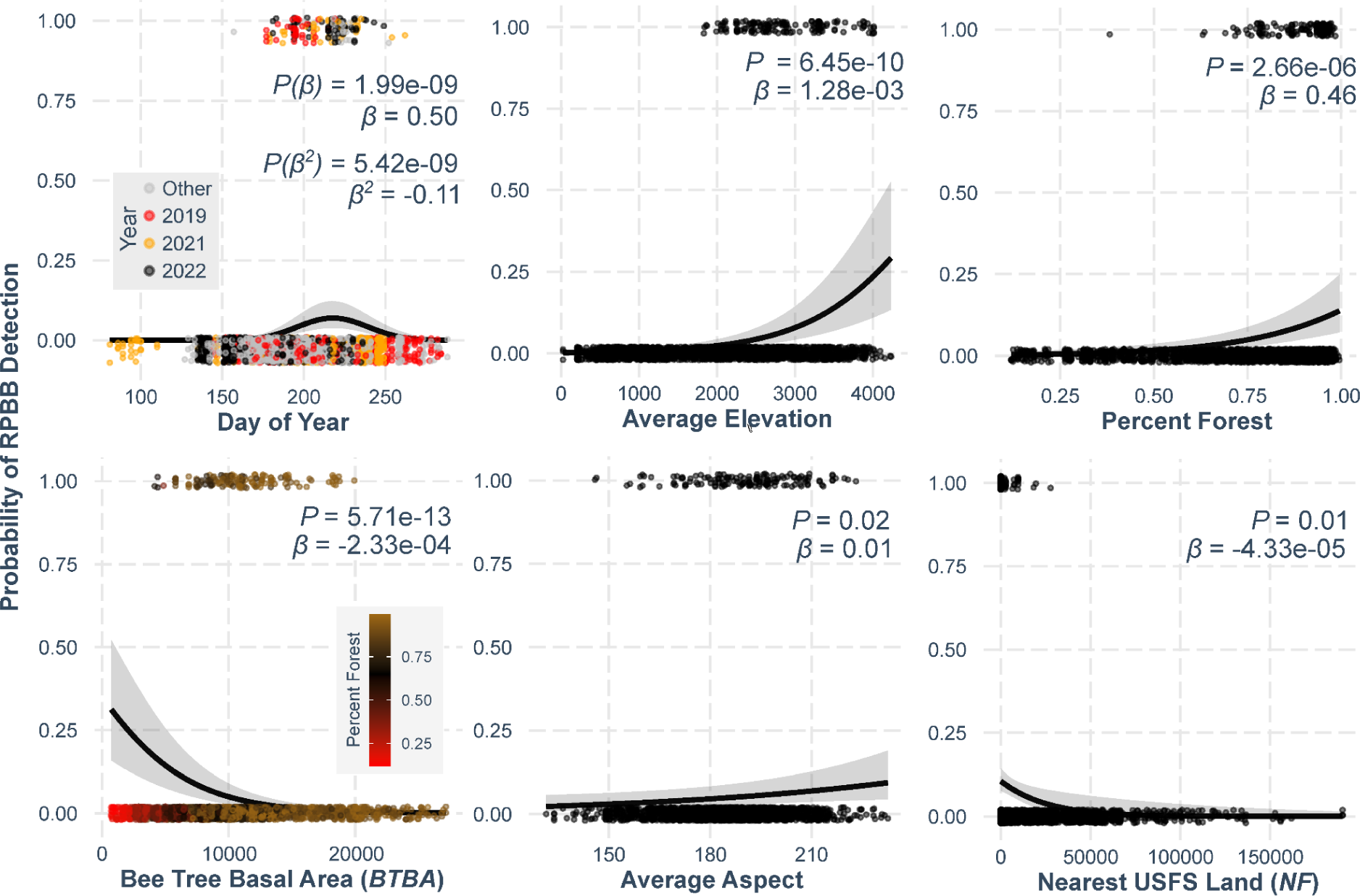
Effect plots of best-fit relationships for the montane-centric model (Equation 1) of RPBB presence-absence (1/0) outcomes. *Forest*, *BTBA* and average *Elevation* and *Aspect* were estimated at a 3000 m radius surrounding each survey point. *Elevation* is plotted in feet, with *NF* in meters. *BTBA* represents square feet of stem basal area per acre. In plotting *Day*, a small y-axis offset was added to facilitate visualization of detections by year. Effect plots were generated assuming an average elevation of 2,600 ft above sea level (i.e. *Elevation* = 2,600) to illustrate expected detection probability in relatively favorable climatic conditions.

Selection for a more generalizable model based on bioclimatic variables (Equation 2, Figure 3) resulted in the substitution of *BIO8* and *BIO9* for *Elevation* and *Aspect*. This substitution considerably improved model fit (McFadden’s pseudo-*R^2^*= 0.379; ROC AUC = 0.919; ΔWAIC = -72.4). Inferred effects were qualitatively and quantitatively similar across the montane and bioclimatic models, as supported by an OLS regression comparing coefficients of terms common to both models (Intercept CI: -0.684 to 1.612, *β* CI: 0.730 to 0.819, *R^2^* = 0.998). Minimal evidence of multi-collinearity between predictors was observed in this final model (VIF < 1.5 for all terms).

**Figure 3:**
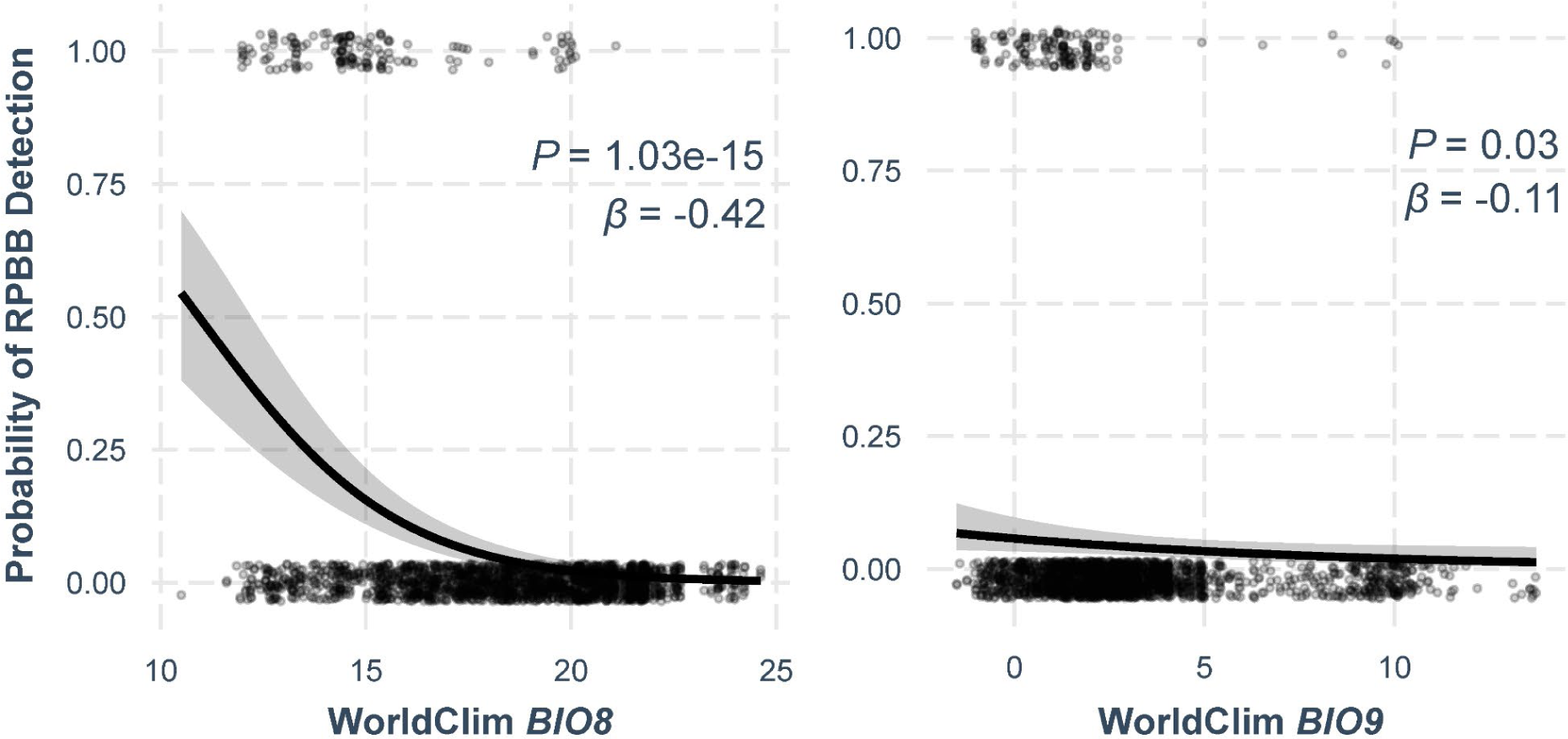
Effect plots of best-fit relationships for the bioclimatic model (Equation 2) of RPBB presence-absence (1/0) outcomes in association with the average *BIO8* and *BIO9* temperatures at a 3000 m scale. Effect plots were generated assuming surveys were conducted on US National Forest lands (i.e. *NF* = 0) to illustrate detection probability in favorable forest habitat conditions.

Incorporating the basal areas of *Pinus* and *Quercus* into the modeling process resulted in the removal of *BTBA, Forest* and *BIO9* during stepwise selection and revealed significant positive associations between RPBB and these genera (*P* < 0.001 for both terms). The final model (Equation 3; Figure 4) was well-fit to the data (McFadden’s pseudo-*R^2^* = 0.379; ROC AUC = 0.919) and there was minimal evidence of multi-collinearity between predictors (VIF < 1.5 for all terms).

**Figure 4:**
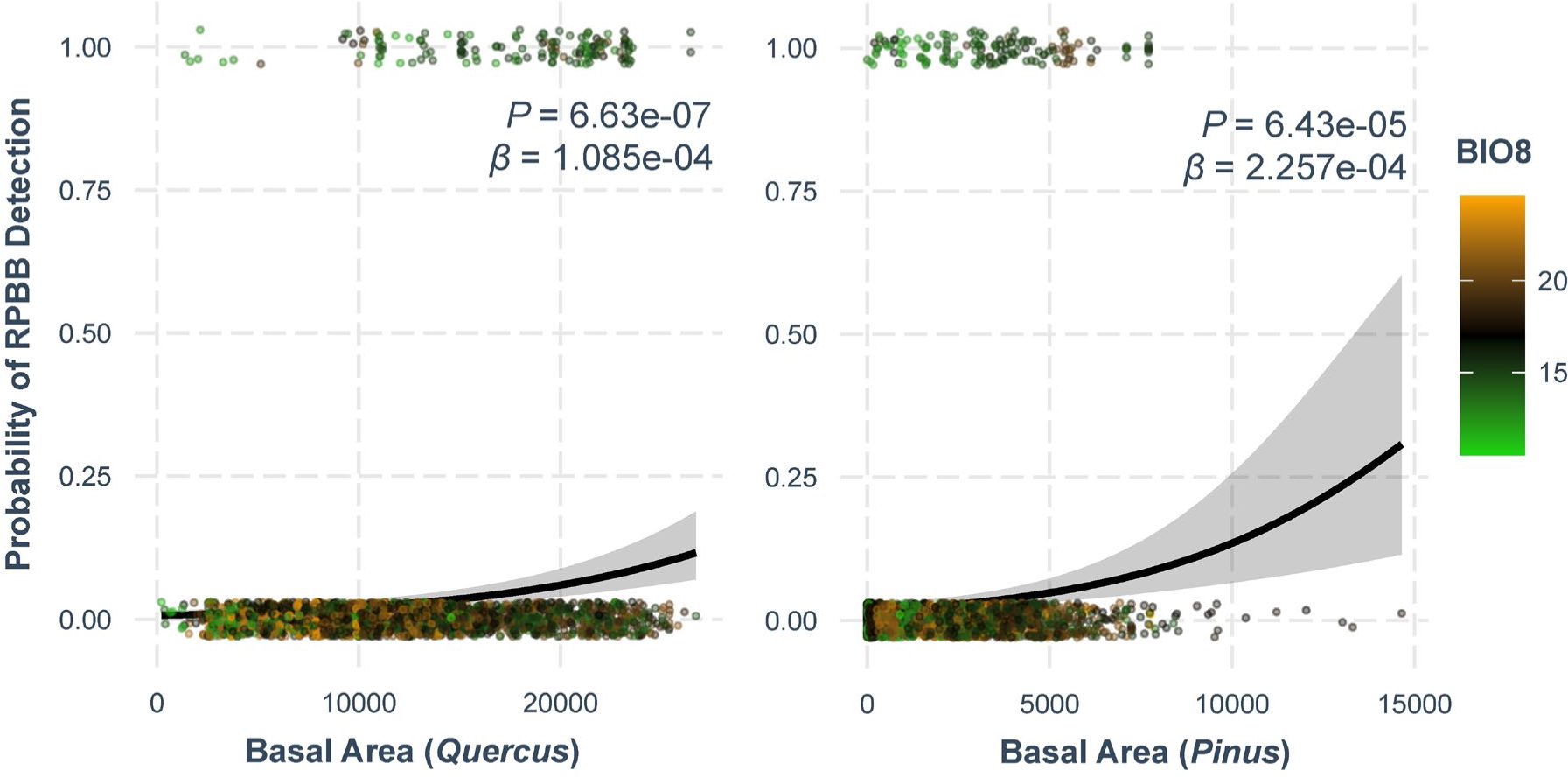
Effect plots of best-fit relationships for the extended tree basal area model (Equation 3) of RPBB presence-absence (1/0) outcomes in association with the basal areas of *Quercus* and *Pinus* at a 3000 m scale.

## Discussion

With these observations, we document a persistent population of RPBB in Central Appalachia based on repeated observations across several years. Analysis of local habitat characteristics suggests that RPBB is supported by specific climate and habitat conditions: Cool, heavily forested, high-elevation landscapes. Therefore, it is not likely to be widespread across the eastern U.S. Through multiple complementary models that account for climatic and seasonal trends, we show that forest composition is strongly associated with RPBB presence. Unexpectedly, after controlling for other variables, surveying in areas with high basal area of putative spring forage tree species (*BTBA*) appears to decrease the likelihood of observing RPBB. In exploring this trend further, it is plausible that the functional explanation for a negative association with *BTBA* is that many of the preferred RPBB plant hosts occur primarily in the understory of non-*BTBA* plant canopies (i.e., oak-pine forest). Within Central Appalachia, mixed forests containing fire-tolerant *Quercus* and *Pinus* stands are common. Such forests are typically associated with understory communities of Ericaceae and Rosaceae, which are potentially important forage species and often contain a high species richness of spring ephemerals. Lastly, though species-level forest composition was not investigated, previous work suggests that forests provide preferred nesting habitat for many bumble bee species (Lanterman et al., 2019; Liczner & Colla, 2019), including RPBB (Boone et al., 2021).

Aside from climatic associations, Central Appalachian landscapes associated with RPBB are typified by abundant native and mass-flowering species. In the spring and early summer, mass blooms of shrubby early successional species (e.g., *Aronia, Rubus* and *Rhus*), mid-story species (e.g., *Amelanchier, Prunus* and *Crataegus*), shrubby Ericaceae (e.g., *Pieris*, *Vaccinium, Gaylussacia, Rhododendron* and *Kalmia*) and canopy species (e.g., *Prunus*, *Tilia*, *Oxydendrum*, *Nyssa* and *Liriodendron*) are abundant. Throughout midsummer and fall, forage items include understory and early successional species (*Hydrangea, Aralia, Robinia* and *Rubus*) as well as field and forest edge associates including various Asteraceae (e.g., *Solidago*, *Eutrochium, Helianthus, Verbesina, Cirsium,* and *Ageratina*), Lamiaceae (*Pycnanthemum*, *Monarda*, *Prunella* and *Agastache*) and other native forbes (e.g., *Actaea* and *Asclepias*).

Among these forage items, a number of native taxa appear to provide preferred late-season pollen and nectar. Previous work suggests floral resources are scarce within forested regions during this time (Proesmans et al., 2019; Mathis et al., 2022; Sponsler and Johnson, 2015) and these species warrant further investigation for their potential to support RPBB populations. Native species such as *Hydrangea arborescens*, *Actaea racemosa*, *Thalictrum pubescens*, *Agastache scrophularifolia*, *Asclepias syriaca*, *Monarda fistulosa*, *Verbesina alternifolia*, *Pycnanthemum incanum*, *Eutrochium fistulosum, Rubus oderata* and *Helianthus divaricatus* would be strong candidates for incorporation into management plans along roadsides and within timberland understories.

While the data suggest forest habitat features are most strongly associated with RPBB, open areas interspersed throughout forested landscapes complement the forage available in Central Appalachian landscapes in important ways. For example, our data includes multiple RPBB detections on non-native forage items within agricultural pastures. These fields provide floral resources in late-summer and altering management of these areas represents an additional avenue toward *Bombus* conservation. Incentives to promote native forb diversity and minimize mowing of fields during late summer and early fall when RPBB are actively foraging at forest edges represents a feasible starting point.

Through intensive surveys of U.S. national forest lands (WV and VA) and state forests (MD and PA), we noted stark differences in roadside and understory vegetation. Roadside and understory habitat within national forest lands are typified by areas of heightened floral diversity and abundance, with numerous patches of preferred host plants residing in close proximity (Hanula et al 2016). These areas of habitat likely provide important resources for RPBB colony establishment, growth and reproductive output. In addition, such habitat facilitates detection of the species within densely forested and rugged montane landscapes. In contrast, we observed few, if any, sites with analogous habitat in the state forest areas we surveyed, which tended to be heavily mowed. In many cases, these differences in understory habitat are apparent despite close proximity and habitat connectivity between state forests and national forests. Differential management practices, including a lack of prescribed fire, frequent roadside mowing, and ditch management may contribute to fewer observations of RPBB within state forests compared to national forests. Additional factors including differential land use history (e.g., differences in the historical intensity of timbering, grazing and burning) may also contribute. Regardless of the causal factors associated with differences in floral resource conditions, management alterations that promote mid to late-summer RPBB forage items could provide additional habitat value while improving our ability to characterize the full spatial extent of RPBB occupancy within north-central Appalachian regions of MD and PA. Initiatives which alter management and floral resources at the edge of the current RPBB range would be particularly useful for gauging the causality of putative associations and potential impacts of implementations within the known RPBB range.

Central Appalachian RPBB appear to forage from similar flower species relative to the Upper Midwest population. Of all plant genera with more than two RPBB floral visitation observations (*n* = 17), eight overlapped with the top 17 most visited genera in the Upper Midwest (Wolf et al., 2022). A number of additional flower genera also overlapped with this study including, but not limited to, *Carduus*, *Physostegia*, *Melilotus*, *Monarda*, *Rubus,* and *Actaea*. Similarly, our data contain RPBB observations from approximately 45 percent of the floral genera on which RPBB was observed in the Upper Midwest (Otto et al., 2023), despite the fact that those surveys were conducted predominantly in urban and agricultural landscapes. Overall, our data support conclusions based on historic observations that RPBB is a highly generalized forager (Colla & Dumesh, 2010), with differences across regions likely reflecting differential availability of floral resources in preferred wetland, prairie, and forested habitats.

The extent to which a mosaic of land-use patterns contributes to the persistence of RPBB within this region warrants further investigation. The use of controlled burns and variable timbering practices (often performed by USFS) plays a clear role in driving floral resource availability of key forage plants in forests with implications for bee forage (Brown et al., 2017; Burkle et al., 2019; Zellweger et al., 2020; Gelles et al., 2022). These disturbances restart forest succession, increasing the reproductive investment of many species (Newell & Tramer, 1978), and contributing to dense floral patches with larger floral rewards (Kudo et al., 2008). Knowledge gaps in our understanding of how forest management influences floral resources and nesting habitat, and via these factors, bee populations themselves, make it difficult to predict which management practices are most beneficial (Rivers et al 2018) and bumble bee species show differential responses to forested land (Novotny et al. 2021). Nevertheless, there is evidence that some management practices aimed at other wildlife goals also benefit bumble bees (Mola et al. 2021). Given the positive association between RPBB occurrence and proximity to USFS lands, exploring the influence of disturbance regimes associated with current USFS management is critical. A study designed to partition the influence of fire history, timbering practices, stand age, and tree species composition on bee forage and RPBB presence would help inform conservation management for this endangered species.

Overall, our research suggests that forest extent and composition is strongly associated with RPBB persistence. Across the Eastern U.S., forests have undergone a well-documented transition from oak-dominated pyrophytic communities to pyrophobic communities (Alexander et al., 2021; Nowacki & Abrams, 2008). This process, termed mesophication, has been documented within the Appalachians (Dyer & Hutchinson, 2019; Flatley et al., 2015) and may have implications for RPBB. Our work shows that RPBB are more likely to be encountered within oak and pine-dominated landscapes. The predominant counterpart, forests composed primarily of beech, maple and birch species (Schroeder et al., 1997; Supplemental Figure S2), appears less associated with RPBB in Appalachia, perhaps due to differences in nesting resources (Clotfelter et al., 2007) or understory species associates. Species associated with forest mesophication generally lead to less flammable understory leaf litter (Kreye et al., 2013), perpetuating a positive feedback loop for further losses of pyrophytic communities and decreases in fire within the landscape. Research on fire dynamics and pollinators suggest that burn severity and extent are associated with broad measures of pollinator abundance and diversity (Galbraith et al., 2019; Ulyshen et al., 2022; Potts et al., 2003), highlighting the need for studies focused on RPBB specifically.

Recent works emphasize that causal drivers of bumble bee decline include climate change and pesticide use as major factors (Janousek et al., 2023; Soroye et al., 2020). While these factors likely play a role in RPBB decline, the persistence of the species in two apparently distinct populations within Central Appalachia and the Upper Midwest is unlikely to be fully explained by these particular anthropogenic forces if the currently known range of the species is relatively accurate. In contrast to the Upper Midwest RPBB population, the extent of agricultural and urban land was not strongly related to RPBB occurrence, though this lack of association may reflect substantially less large-scale agriculture and developed land cover in the region studied (Boone et al. 2023). Furthermore, if climate change and pesticides were the predominant drivers of decline, the presence of the species in the Upper Midwest and Central Appalachia, along with the absence of the species in Indiana, Michigan, Southern Ontario, and the Northeast, would be unexpected. Our data suggest that certain areas of Indiana, Michigan, Southern Ontario and the Northeast would provide favorable climatic conditions for the RPBB. Within these areas, pesticide exposure risks are variable and often considerable (Krupke et al., 2012; Shaafsma et al., 2015), but there are relatively low risk areas that are not likely to exceed the exposure risks experienced by extant populations in the Upper Midwest (Douglas et al., 2020). Additionally, the recent work by Jackson et al. (2022) highlights that inferred associations between species occupancy and anthropogenic risk factors are highly species-specific. In the case of members of the subgenus *Bombus*, population declines occurred rapidly and coincided with exceptional rates of pathogenic infection and load (Cameron et al., 2011; Colla & Packer, 2008; Grixti et al., 2009). This association suggests that climate, habitat degradation, and pesticide risk were more likely to have played interacting roles with some other rapidly acting causal driver such as a swiftly spreading parasite. Under this scenario, rapid recolonization and dispersal of the species may be limited by habitat fragmentation in many areas, since successful recolonization is rare within a matrix of small and isolated patches of favorable habitat (Bowers et al., 1985).

Based on floral visitations observed in this study, the role of highly abundant and mass flowering species with unique secondary chemistry is worth investigating. Flowers from the families Ericaceae, Apocynaceae, and Ranunculaceae, along with certain Asteraceae, are known to produce nectar and pollen containing secondary metabolites including cardenolides, diterpenoids, and glucosides with strong bioactive potential (Adler, 2000; Agrawal et al., 2012; Rivest & Forrest, 2020; Stevenson et al., 2017). Generally, bumble bees appear adapted to consuming nutritionally relevant doses of these compounds (Jones et al., 2023), and their presence in the bumble bee diet may have implications for behavior (Villalona et al., 2020) and the epidemiology of common trypanosomal and microsporidial pathogens (Bernklau et al., 2019; Fowler et al., 2022; Palmer-Young et al., 2017; Richardson et al., 2015). Interestingly, recent work in the Appalachian ridge and valley physiographic region reported variation among habitats in the incidence of bumble bee pathogens, with trypanosome infections higher and viral infections lower in ridgetops relative to valleys (Gratton et al., 2023). The underlying drivers of this association remain unexplored. Regardless of the explanation, positive associations between trypanosomal infections and overwintering survival have been documented in *Apis mellifera* (Richardson et al., 2023), and further studies are needed to test whether this association can be generalized across other Apidae, including *Bombus*.

In using GLMs, we note that our statistical approach does not attempt to separate the detection process from inference of occurrence patterns. An additional consideration is that, while common across ecology and environmental science, inferences from regression models of field observation data can be limited by a number of statistical phenomena (e.g. omitted variable bias, collider bias, multi-collinearity) and relationships should not necessarily be viewed as causal (Arif & MacNeil, 2023). Overall, we show that there are strong relationships between RPBB detection and various landscape features but acknowledge the caveat that predictors can be related to each other, as well as to other unmeasured variables within the study system.

Interpretations will be most robust when limited to inferential questions of what landscape characteristics are broadly associated with RPBB, whether directly or indirectly, and predictive questions of where RPBB occurs *and* is relatively more detectable. While the use of an occupancy modeling approach within a highly temporally replicated design may be ideal for certain research questions (Boone et al., 2023; MacKenzie et al., 2006; Otto et al., 2023), it requires substantially greater survey effort to be implemented at the scale we present here. Additionally, the degree to which occupancy modeling assumptions are met with long-distance and non-random foraging habits of bumble bees situated within relatively segregated and patchily-distributed foraging conditions remains poorly explored.

In terms of selection of features, choosing methods for model selection is a strongly contested area within ecological research. Our primary approach involved gathering consensus across five independent research teams to agree on an initial model and hypotheses, followed by reverse stepwise selection. However, we also employed a machine learning method, random forest, which relies on a more automated ensemble-based approach to selecting features and modelling associations. Despite typical advantages with respect to predictive accuracy, random forest is generally less interpretable relative to traditional regression modelling and it lacks a framework for probabilistic inferences on individual features. Regardless, random forest models suggested similar associations between landscape variables and RPBB detections when applied to our data (Supplemental Figure S3).

With regard to future survey efforts, our findings suggest that montane areas of PA, North Carolina (NC) and Tennessee (TN) provide appropriate environmental conditions for RPBB persistence in addition to the areas we surveyed. In the absence of adequate, robust survey effort, there is still uncertainty about the long-term persistence of RPBB within these Appalachian regions. At present, observations of Yellow-Banded Bumble Bee (*B. terricola*; YBBB), a closely related generalist species within the subgenus *Bombus*, may represent the best predictor of high quality RPBB habitat. YBBB, which has experienced a range contraction into high elevation areas in the northeast (Jacobson et al. 2018), has also been observed throughout the Central Appalachian RPBB range (Hepner & Smith, 2019) and more recently there have been multiple observations across Western MD, South-Central PA and Western NC (personal observations, RTR and MJH; inaturalist.org). Determining the distribution of bumble bee species of conservation concern within these regions is ongoing and it is unknown whether these detections reflect increases in population size or simply increased survey effort. Since RPBB detection is highly dependent on surveying appropriate habitat within narrow time intervals, we feel that considerably more survey effort is needed to ensure a more complete understanding of the extent and health of RPBB populations within rugged, forested, and montane areas of the Eastern U.S. and throughout U.S. Fish and Wildlife Service conservation units four and five. Such information will be critical for guiding conservation management, as well as consideration of future de-listing of the species (U.S. Fish and Wildlife Service, 2021).

## Author Contributions

TR and RPJ were responsible for the initial discovery of Central Appalachian RPBB populations. MJH, RPJ, EO, KP, TR and RTR conducted surveys and provided RPBB sightings for analysis. RTR, MJH, TR, KG, KP, and RPJ conceived and designed the analysis. RTR conducted all statistical analysis and visualization. RTR, MJH, KG, RPJ and TR wrote the initial draft of the manuscript. All authors contributed to the editing and reviewing of the manuscript.

## Acknowledgements

These surveys were supported by funds from Metamorphic Ecological Research and Consulting, LLC, USFWS Region 5 Science Applications, USFWS -VA Ecological Service, USFWS Section 6 funds administered by MD DNR Natural Heritage Program to MJH; USFWS Section 6 funds administered by VA Department of Agriculture and Consumer Services to EO; funds and resources from Environmental Solutions & Innovations, Inc. to RPJ and KP; and a DoD ESTCP Grant to RTR and KG (Project RC22-B5-7373). All required permits and authorizations were obtained to complete these surveys. We thank USFWS, USFS, Monongahela National Forest, George Washington and Jefferson National Forest, VA DCR-DNH, MD Forest Service, MD Park Service, MD DNR, WV DNR, WV State Parks, The Nature Conservancy - WV, Friends of Deckers Creek, WV Land Trust, WVU Jackson’s Mill, Riffle Farms, and private landowners who aided with site access and permitting. On behalf of MJH, we thank family members and colleagues who provided support and guidance, particularly Walter Veselka. The authors thank Matthew Fitzpatrick for aggregation of survey site landscape statistics. Additional thanks to Michelle Jean, Andrew LaVoie, Paige Reeher, Doug Gilbert, Grace Avalos and Cameron Garland for additional help in the field locating RPBB, and to Allstar Ecology, LLC for use of survey equipment in 2018-2019.

## Supplemental Material

**Supplemental Figure S1:**
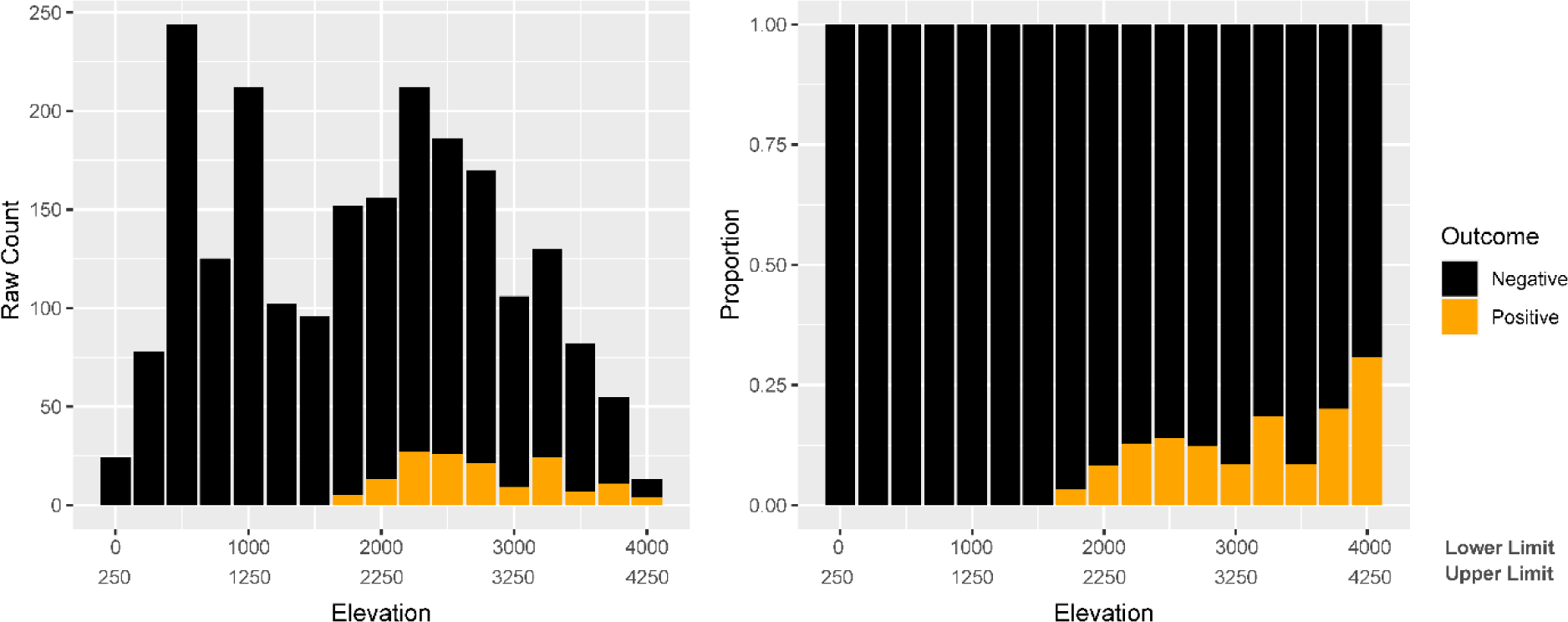
Illustration of the elevations at which RPBB were observed in our data. For this analysis, surveys were grouped into bins spanning 250 ft ranges of average elevation. Data are shown by raw count and proportion.

**Supplemental Figure S2:**
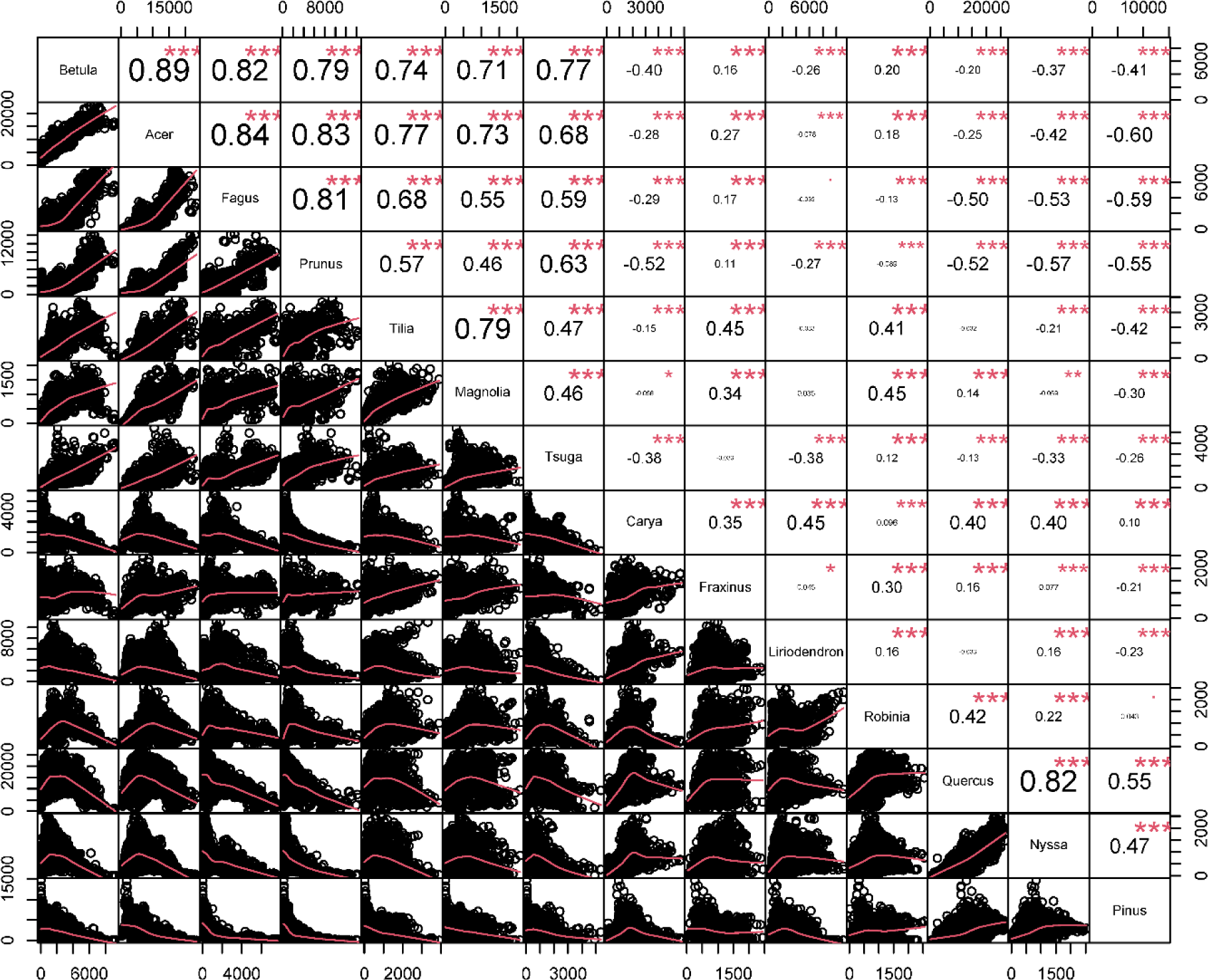
Illustration of associations between dominant tree genera, those with cumulative basal area greater than one million square feet per acre, across all study sites (measured at a 3 km radius). Correlations illustrate the predominance of two major forest types; 1) beech-maple-birch forests which are typically mixed with associates including cherry and basswood, and 2) oak-pine forests, which tend to be associated with black tupelo (*Nyssa sylvatica*).

**Supplemental Figure S3:**
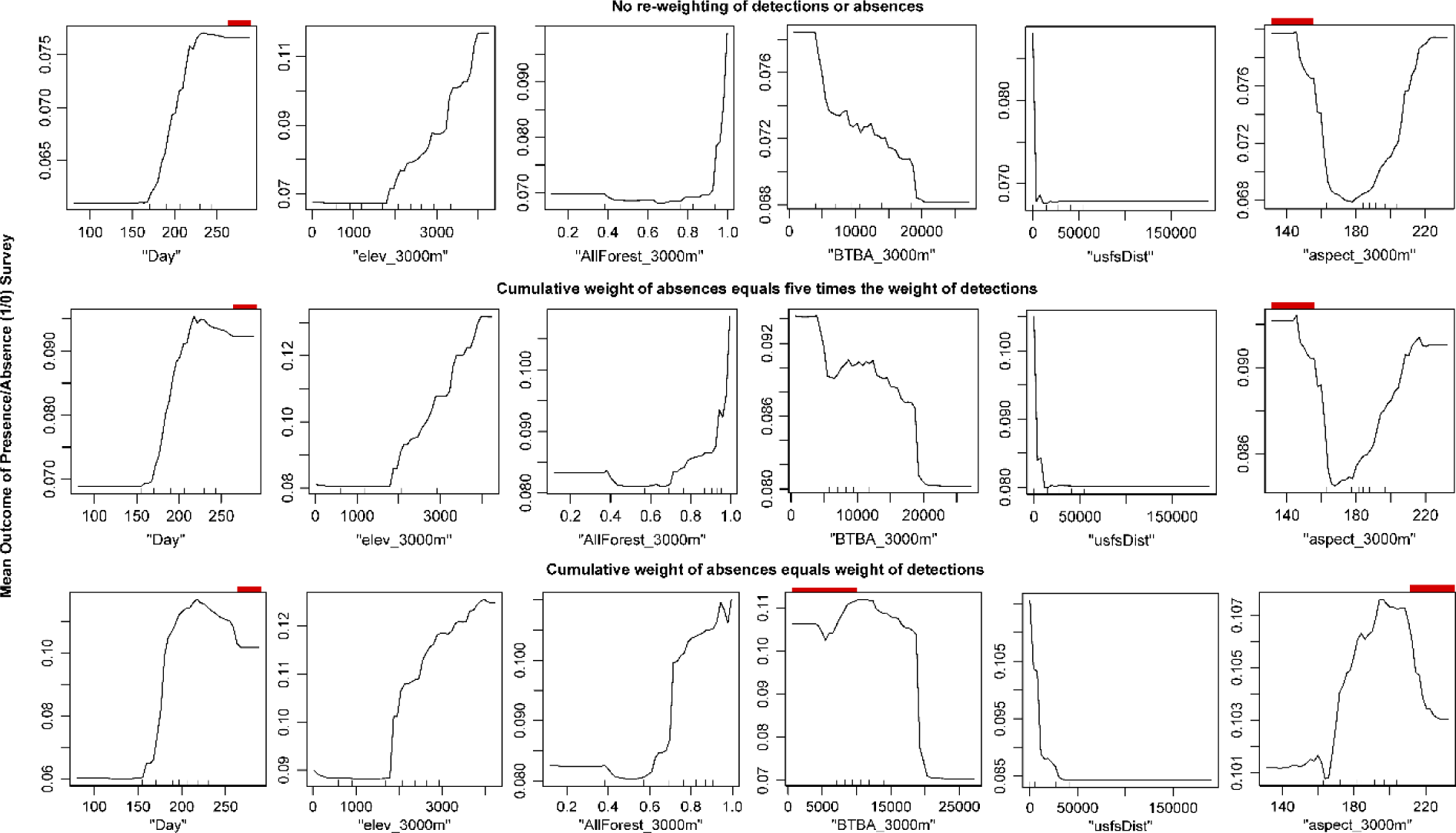
Random forest partial dependence plots for the variables included in GLM Equation 1. During random forest regression, RPBB survey presence and absence outcomes were regressed against 42 predictor variables including landscape composition and topography measures, bioclimatic variables and the day of year of the survey. Three distinct sets of weights were applied to evaluate the robustness of results at multiple levels of down-weighting survey absences, which were about 20 times more prevalent in the data relative to detections. Horizontal ticks along the x-axis show the deciles of the distribution for each predictor variable. Plots suggest that random forest interpretations are largely congruent with those of GLMs, with few exceptions. Red bars highlight areas of the predictor space where the association is incongruent with GLMs. It should be noted that, in many cases, strongly non-monotonic effects are inferred based on relatively small numbers of data points using the random forest method.

**Supplemental Table S1:** Environmental variables used for statistical analysis, each of which was calculated at a 500 m, 1000 m and 3000 m radius surrounding each survey site.

| Source | Variable |
| --- | --- |
| USDA Forest Service | usfsDist |
| Daymet | bio1 |
|  | bio2 |
|  | bio3 |
|  | bio4 |
|  | bio5 |
|  | bio6 |
|  | bio7 |
|  | bio8 |
|  | bio9 |
|  | bio10 |
|  | bio11 |
|  | bio12 |
|  | bio13 |
|  | bio14 |
|  | bio15 |
|  | bio16 |
|  | bio17 |
|  | bio18 |
|  | bio19 |
| Open Topography Global Dataset | elevation |
|  | aspect |
|  | slope |
|  | tri (terrain ruggedness) |
|  | roughness |
| National Land Cover Dataset | water |
|  | dev_openSpace |
|  | dev_low |
|  | dev_med |
|  | dev_high |
|  | barren |
|  | mixed_forest |
|  | deciduous_forest |
|  | evergreen_forest |
|  | shrub_scrub |
|  | grassland_herbaceous |
|  | pasture_hay |
|  | cultivated_crops |
|  | woody_wetlands |
|  | emergent_herb_wetlands |
| Forest Inventory Analysis | BA_allTreeSpp |
|  | BTBA |

**Supplemental Table S2:**
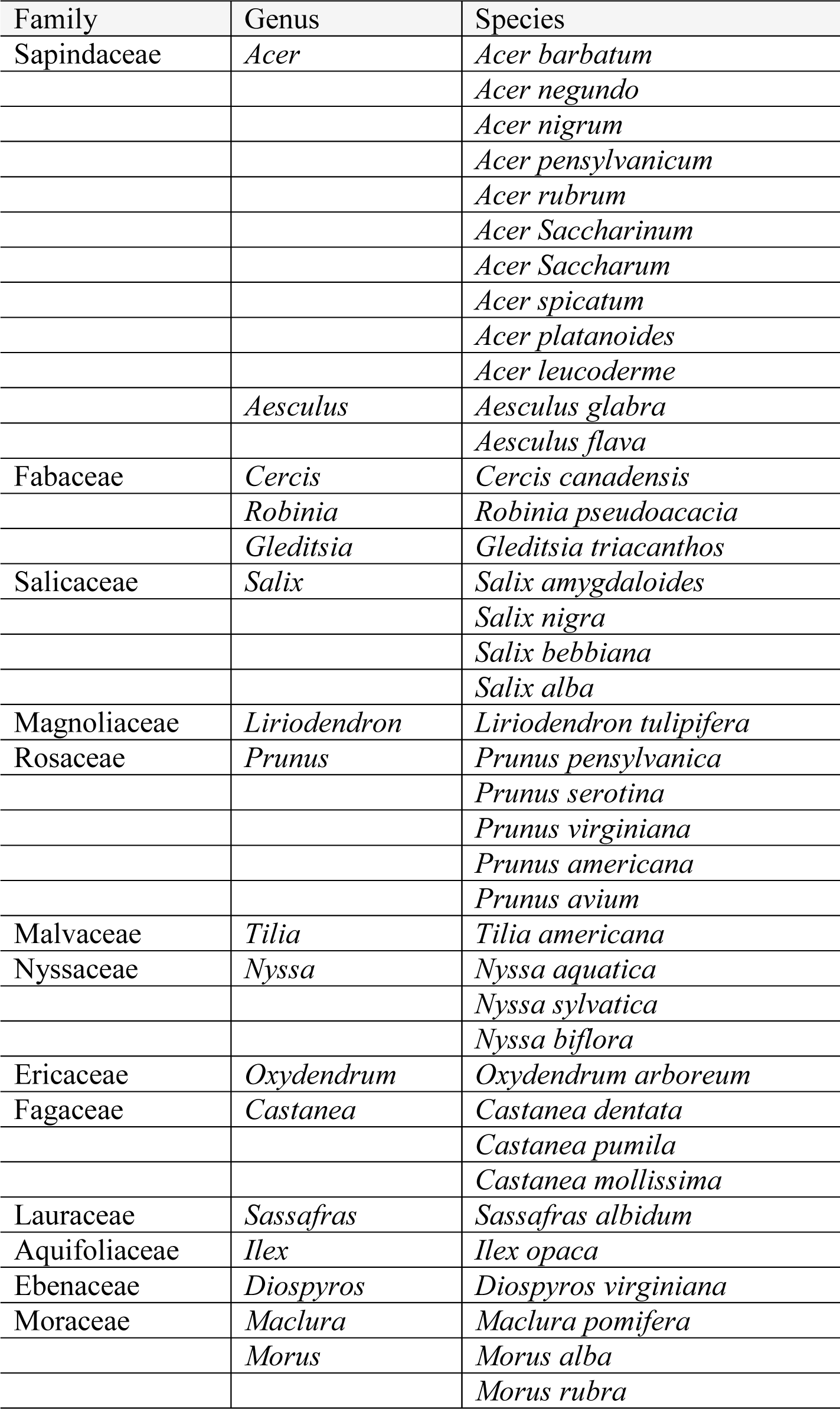
List of ‘bee friendly’ flowering tree species (*BTBA*).

## Notes

### Competing Interest Statement

The authors have declared no competing interest.

